# Chewing phase - theta amplitude coupling facilitates working memory

**DOI:** 10.1101/2025.11.17.688859

**Authors:** Sebastian Espinoza, Wael El-Deredy, Daniel Moraga-Espinoza, Luis Morreal-Ortega

## Abstract

Rhythmic chewing enhances cognitive performance, yet the neural mechanisms linking oromotor behavior to cortical dynamics remain poorly understood, in part due to motion artifacts that obscure brain signals. We used high-density EEG with a custom artifact attenuation pipeline to examine how mastication shapes frontocentral theta oscillations during working memory.Thirty-one participants performed a visuospatial 2-back task under two conditions: at rest and while chewing. Chewing resulted in faster responses and a selective increase in theta power (4–7 Hz) during the late post-stimulus window (900–1300 ms), a phase associated with cognitive control. Theta amplitude was modulated by the chewing phase, revealing strong cross-frequency coupling between motor output and neural oscillations. The strength of this entrainment increased with chewing frequency, indicating a dose-dependent neural gain. Our results demonstrate that peripheral rhythmic activity can synchronize brain rhythms relevant to cognition in real time, supporting a mechanistic link between bodily action and executive function. These findings enhance our understanding of sensorimotor–cognitive integration and point to natural motor rhythms as promising non-invasive tools for modulating brain activity.

## Introduction

Chewing, a fundamental sensorimotor rhythm, has also been shown to modulate cognitive function. Prior studies have demonstrated that mastication affects attention, working memory, and executive control^1–4^. Gum-chewing experiments have reported improved reaction times, alertness, and task-switching, especially under high cognitive load^5–7^. These effects likely reflect enhanced engagement of neural systems supporting sustained attention and cognitive control. Neuroimaging studies support this view, showing increased activation in the prefrontal cortex, cingulate gyrus, and thalamus during chewing^8,9^. These findings highlight the involvement of cortico-subcortical circuits in mastication-related cognitive effects. However, current findings still fail to clarify how chewing shapes cortical dynamics at the neural level.

Mastication recruits a distributed sensorimotor network, including the brainstem, thalamus, somatosensory cortex, and prefrontal areas^10,11^. However, the specific neural mechanisms through which chewing modulates cognition remain unclear. Most studies apply offline designs, measuring brain activity before or after chewing episodes^7–9^, thereby failing to capture the real-time interplay between chewing and cortical dynamics. The use of diverse methodologies—from fNIRS and EEG to animal models—further fragments the literature. For instance, fNIRS studies report increased prefrontal activation during mastication^8,9^, while rodent models show that chewing promotes hippocampal synaptic plasticity and cognitive resilience^2,12^. Other studies, however, fail to detect consistent cognitive benefits of chewing, likely due to methodological variability or insufficient control of chewing-related artifacts in neurophysiological recordings^13^. These inconsistencies highlight the need for methods that disentangle real-time brain activity from chewing-related interference.

Mastication is thought to exert its neuromodulatory effects via afferent signals from periodontal and orofacial mechanoreceptors. These sensory inputs project to the brainstem and thalamus, activating sensorimotor and associative cortical areas^14,15^. Such afferent-driven activity may transiently enhance thalamocortical excitability and shift the phase of ongoing cortical oscillations, particularly in frontal regions implicated in attention and executive control^16^. In this way, chewing may act as a peripheral entrainment mechanism that modulates cortical responsiveness during cognitive engagement.

Among cortical rhythms, frontocentral *θ*-band oscillations (4–8 Hz) play a crucial role in cognitive control, working memory maintenance, and top-down attention^17,18^. These oscillations support communication between prefrontal and parietal regions and increase power during cognitively demanding tasks. Because *θ* rhythms are functionally relevant and spectrally close to chewing-related EMG activity, they provide an optimal window into the dynamic interaction between mastication and cortical processing.

EEG captures cortical activity with high temporal resolution, but mastication introduces high-amplitude muscle artifacts that significantly challenge signal interpretation. Jaw and facial muscle activities generate broadband signals that obscure brain oscillations, particularly in the *θ* range^19,20^. These artifacts overlap with neural signals in frequency and time domains, complicating analysis with conventional preprocessing methods^21,22^. The absence of artifact-robust pipelines for EEG acquired during chewing has largely hindered real-time investigations of neurocognitive activity. Recent studies emphasize the urgency of developing artifact attenuation techniques capable of recovering valid neural signals under these conditions^23–25^.

To overcome the methodological challenges of chewing-related artifacts, we developed and validated an EEG preprocessing pipeline designed explicitly for naturalistic mastication (See Supplementary Figure S1 for details). This approach integrates Artifact Subspace Reconstruction (ASR)^26^, a pseudo-reference technique adapted from iCanClean^20^, to isolate transient EMG bursts while preserving physiologically meaningful neural signals^21^, and Independent Component Analysis (ICA). Our method more accurately preserved cortical *θ* oscillations in both spectral and temporal domains than conventional pipelines, enabling artifact-resistant EEG analysis and high-resolution monitoring of neural activity during sensorimotor tasks.

We then applied this pipeline to investigate how chewing rhythm modulates cortical activity during a visuospatial working memory, which is represented by *θ* oscillation^27,28^. Specifically, we assessed whether the phase of the chewing cycle modulates *θ*-band amplitude (4–8 Hz) in frontocentral EEG electrodes through phase-amplitude coupling (PAC). This dynamic analysis allowed us to test whether chewing entrains brain oscillations in real time. Our findings reveal that chewing not only improves cognitive performance but also modulates brain rhythms via cross-frequency coupling, providing novel evidence for sensorimotor–cognitive integration during natural behavior.

## Results

Thirty-one participants completed a previously reported 2-back working memory task^29^ in two blocks, with the second block performed while chewing a neutral-flavored gum. To control for practice effects, a control group (n = 15) performed the same task, repeating the 2nd block without chewing.

### Chewing during task speeds up reaction time

In the 2-Back task, the control group exhibited a median reaction time (RT) of 579 ± 27.7 ms (mean absolute deviation, MAD) with an accuracy rate (ACC) of 90.3% during the first block. In the second block, their RT improved to 547.9 ± 46.4 ms, with a slight drop in accuracy to 88.2%. The experimental (chewing) group showed an RT of 617.9 ± 43.1 ms and an ACC of 90.3% in the non-chewing block. However, during the chewing block, RT significantly improved to 543.7 ± 38.8 ms, accompanied by an increase in ACC to 93.4%. This represents a 13.4% reduction in RT for the chewing group, compared to 6% in the control group—a statistically significant difference (p = 0.0321, see Figure 1, panel A).

**Figure 1.**
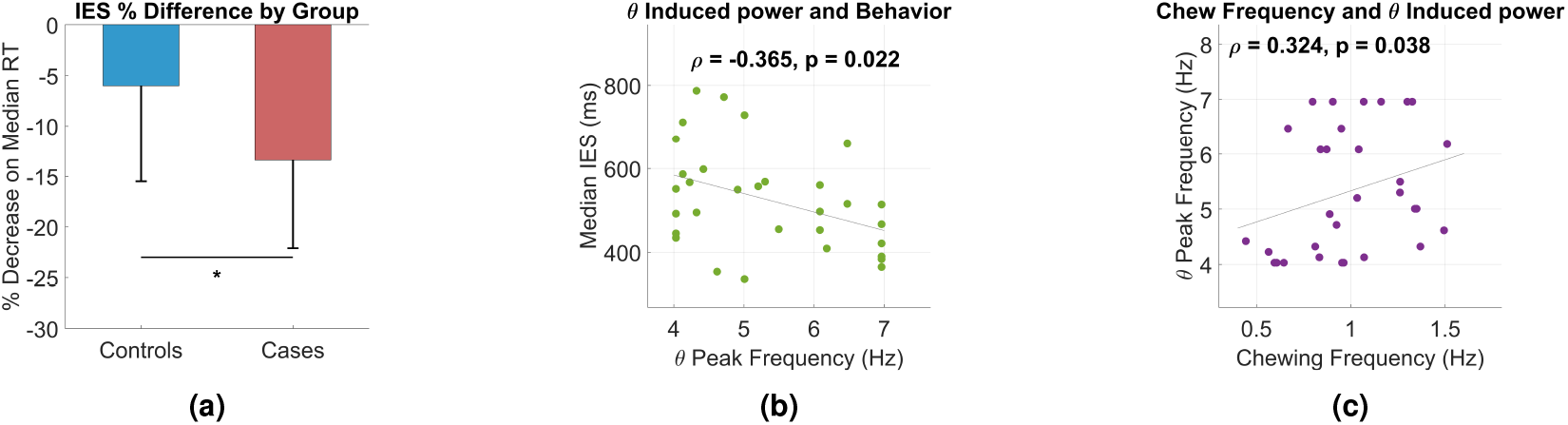
Chewing reduces reaction time during the 2-back task, an effect correlated with increased frontocentral *θ* power. **a** Percentage difference of the inverse efficiency score between subjects who chew and those who do not.**b** Spearman’s correlation showing that as IES decreases, the *θ* peak ([4–7] Hz, [900–1300] ms) increases. **c** *θ* peak increases as chewing frequency increases.

### Frontocentral *θ* reflects changes in the visuospatial working memory processing

Clean, artifact-free EEG trials were analyzed using a Morlet wavelet with a logarithmic increase from 4 to 13 cycles, depending on the frequency. Due to the task’s difficulty, only trials with correct responses were used. Over 4,240 correct trials were used, comprising 2,051 from block 1 and 2,189 from block 2. From each subject we obtain 64.4 ± 17.7 trials (block 1 = 66.2 ±18.7; block 2 70.6 ± 16.7)

Behavioral performance was mirrored in the EEG *θ* frequency band (4–7 Hz), where participants who chewed during the second block exhibited increased *θ* power and reduced inverse efficiency score (IES) between 900–1300 ms post-stimulus (Spearman’s *ρ* =-.3654; p = 0.0216. Figure 1, panel B.). These neural changes are indicative of more efficient cognitive processing. Chewing frequency positively correlated with *θ* peak amplitude (Spearman’s *ρ* = 0.3242; p = 0.0376; Figure 1, panel C.), suggesting a dose–response relationship between rhythmic Chewing and *θ*-band enhancement.

Frontocentral *θ* reflects Changes in Visuospatial Working Memory Processing During the early post-stimulus window (100–900 ms). No significant differences in *θ* power amplitude were observed between the chewing and non-chewing blocks (p = 0.095). However, in the later time window (900–1300 ms), the control group exhibited a significant decrease in induced *θ* power (p = 0.041. See Figure S3), whereas the case group showed a recovery and further increase in *θ* power during the chewing condition(p = 0.005, Figure 2).

**Figure 2.**
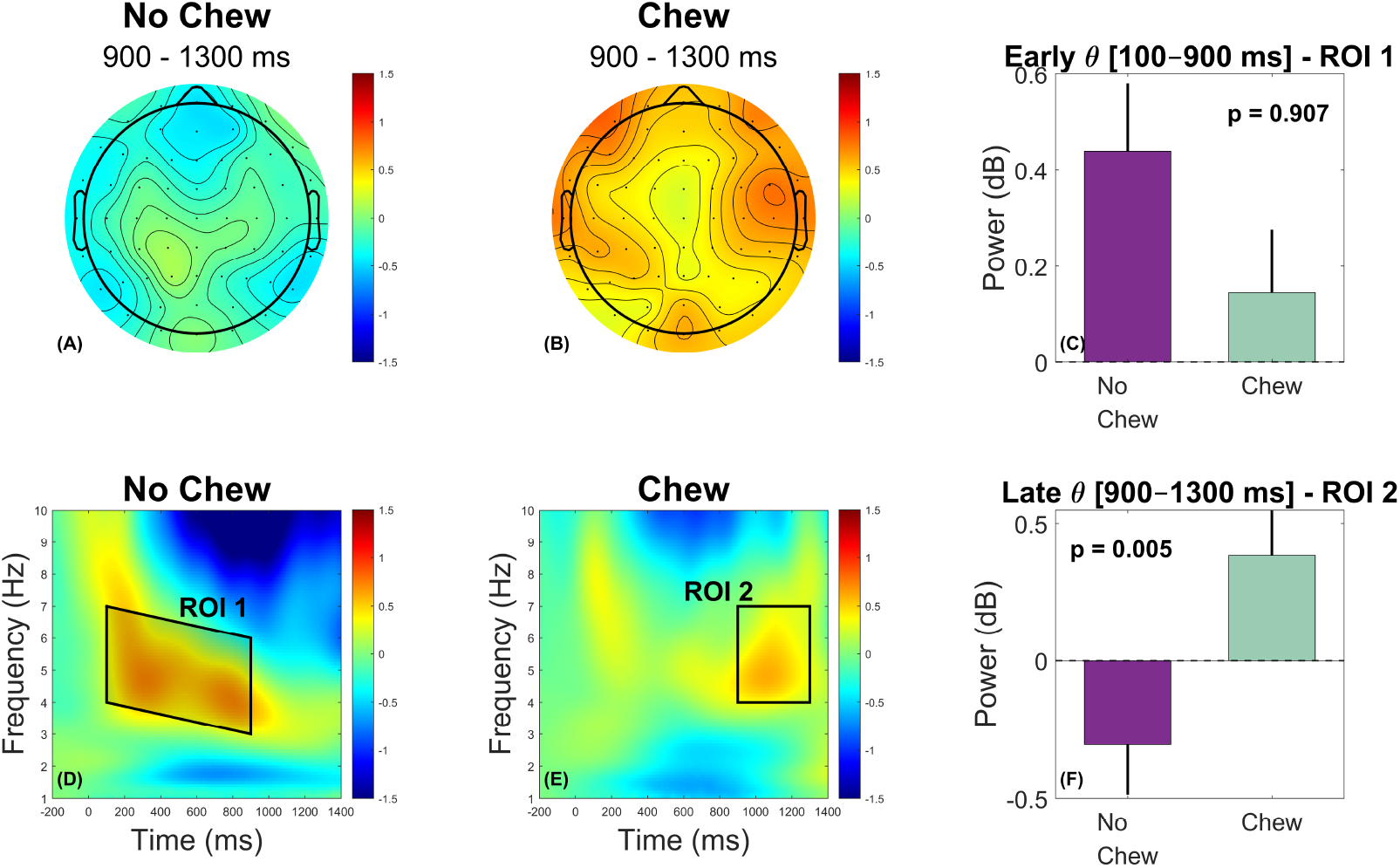
Frontocentral *θ* oscillatory dynamics differ between the early and late phases of the trial, reflecting evolving cognitive control demands. Early in the trial (Panel **D**, ROI 1), *θ* oscillations increase (Panel **C**) in the non-chewing blocks. Late in the trials (Panel **E**, ROI 2), *θ* oscillations increase (Panel **F**) in the chewing blocks. The topographical maps show the frontocentral increase in the late *θ*.

### Chewing rhythm entrains *θ*-band amplitude

To investigate cross-frequency coupling between chewing rhythm and cortical oscillations, we performed a phase–amplitude coupling (PAC) analysis. Specifically, we examined the coupling between the low-frequency phase of the EMG signal (1.0 Hz) and the amplitude of *θ*-band activity (6.5 Hz), averaged across frontal EEG channels—the phase–frequency pair that maximized the group-level modulation index (MI). Figure 3, Panel C, shows a polar representation of the preferred PAC phase across participants. Rather than a uniform distribution, subjects clustered around a consistent phase angle (mean direction = 31.76°; resultant vector length *R* = 0.405). The Rayleigh test confirmed significant non-uniformity (*Z* = 5.097, *p* = 0.0053), indicating a reliable group-level preferred phase of maximal *θ* amplitude modulation during the chewing cycle.

**Figure 3.**
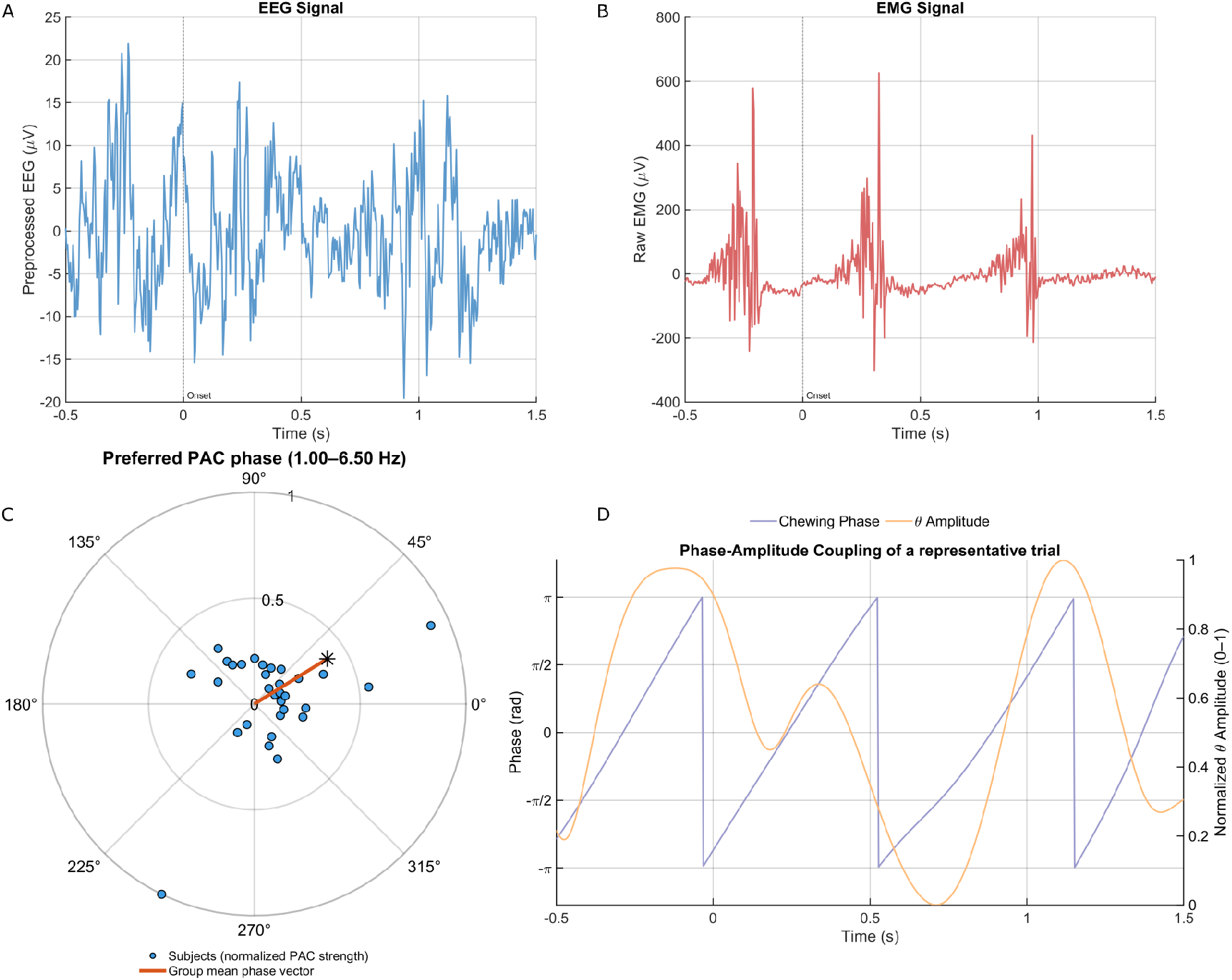
Chewing rhythmically modulates *θ* amplitude through phase–amplitude coupling, linking motor output to neural oscillations. Across the continuous data, the phase of the chewing signal modulates the amplitude of the *θ* band. Panel **C** shows the preferred PAC phase across subjects using a polar representation: each point denotes an individual subject’s preferred phase (normalized PAC strength), and the red vector represents the group mean resultant direction at the phase–frequency pair yielding the highest modulation index (MI). Example of 2 seconds of preprocessed EEG from averaged frontocentral electrodes (Panel **A**). Same 2 seconds of raw EMG data from the right masseter muscle (Panel **B**). Chewing phase and *θ* amplitude over the same time window (Panel **D**).

This relationship is illustrated in Figure 3, Panel D, for a representative trial. Fluctuations in *θ* amplitude (yellow) closely follow specific phases of the chewing signal (purple), demonstrating how the masticatory cycle dynamically modulates cortical activity.

At the group level, Tort’s modulation index (MI)^30^ was calculated for each participant. A one-sample Wilcoxon signed-rank test showed that MI values were significantly greater than zero (median MI = 1.40 × 10^−4^; *p <* 0.0001), indicating robust PAC across the cohort. Additionally, 6 out of 31 participants (19.35%) exhibited individually significant MI values (*p <* 0.05), exceeding the rate expected by chance (binomial test, *p* = 0.0039). Together, these results provide strong evidence that *θ*-band cortical activity is systematically modulated by the phase of chewing, reflecting non-random cross-frequency coupling at both individual and group levels.

## Discussion

The present study demonstrates that rhythmic mastication modulates cognitive performance and reorganizes cortical oscillatory dynamics during visuospatial working memory. Using an on-line chewing manipulation, we show that the phase of the masticatory rhythm (0.5–2 Hz) entrains frontocentral *θ*-band activity through phase–amplitude coupling (PAC), and that this entrainment scales with chewing frequency. Critically, this neural modulation co-occurs with behavioral improvements, including faster responses and reduced inverse efficiency scores. These findings identify chewing as a potent source of peripheral rhythmic input that interacts with central executive networks in real time.

Importantly, these results converge with an independent dataset using a post-chewing manipulation, where chewing performed *before* the task similarly improved performance and increased frontocentral *θ* power alongside enhanced frontoparietal connectivity. The temporal locus of the manipulation differed between experiments—online versus pre-task chewing—but both produced the same core signature: a chewing-related facilitation of executive performance coupled with upregulation of frontocentral *θ*. This cross-paradigm convergence substantially strengthens the interpretation that mastication engages a reproducible sensorimotor mechanism for cognitive enhancement rather than producing task-specific or pipeline-specific artifacts.

### Chewing as a bottom-up modulator of cortical *θ*

Frontocentral *θ* oscillations (4–8 Hz) are strongly implicated in performance monitoring, cognitive control, and working memory^17,18^. Our PAC analyses reveal that chewing rhythm modulates the amplitude of *θ* activity, with entrainment emerging predominantly in the late post-stimulus interval (900–1300 ms). This time window is classically associated with response evaluation and top-down control rather than perceptual encoding, suggesting that chewing influences executive rather than sensory stages of information processing. The positive association between chewing frequency and *θ* amplitude further supports a dose–response relationship between sensorimotor input and cortical entrainment.

These findings align with a growing literature demonstrating that slow bodily rhythms—breathing, locomotion, cardiac cycles—modulate faster cortical oscillations through nested cross-frequency coupling^31,32^. Peripheral rhythms create temporal “windows of opportunity” for neural excitability and information flow^33^. Chewing appears to operate within this same class of bodily entrainers, dynamically aligning executive computations with predictable rhythmic structure.

A key methodological contribution of this study is the ability to quantify PAC during active mastication. Chewing introduces intense EMG activity near frontal electrodes, and prior work lacked the capacity to reliably dissociate muscle from cortical signals using EEG^23,24^. Through a conservative preprocessing pipeline (ASR + ICA + iCanClean pseudo-referencing) and explicit validation of EMG–EEG separability, we ensure that the observed *θ* modulation reflects neural rather than muscular sources. This represents an important advance for studying cognition under ecologically valid conditions.

### Mechanistic pathways: thalamocortical resonance and neuromodulatory gain

While PAC and *θ* enhancement provide a robust mesoscale signature of chewing-related modulation, the underlying mechanisms remain partly inferential. Periodic afferent input from orofacial mechanoreceptors reaches trigeminal nuclei, brainstem centers, and multiple thalamic relays^15,34^. The thalamus, in turn, is a resonant hub capable of gating sensory information and synchronizing cortical oscillations through frequency-specific loops with prefrontal and parietal cortices^35,36^. Rhythmic trigeminal input may thus entrain thalamocortical circuits, periodically enhancing *θ* alignment and facilitating top-down control signals.

In parallel, neuromodulatory systems—particularly the locus coeruleus (LC)—may contribute to the observed effects. The LC–noradrenergic system dynamically adjusts cortical gain and attentional precision^37,38^, and is sensitive to rhythmic somatosensory input^39^. Chewing may therefore increase the precision of task-relevant signals through coordinated effects on thalamic resonance and LC-mediated gain modulation. Although the present EEG design cannot directly index deep structures, the joint presence of PAC, *θ* enhancement, and behavioral facilitation is compatible with such multi-level sensorimotor–neuromodulatory interactions.

### Convergence with long-term plasticity mechanisms

Beyond immediate entrainment, chewing has been linked to long-term changes in hippocampal and prefrontal physiology. Animal studies show that masticatory deprivation reduces brain-derived neurotrophic factor (BDNF), impairs long-term potentiation (LTP), and degrades hippocampal–prefrontal *θ* coherence^**?**,**?**,2^. Restoring mastication reverses these effects and rescues memory performance^40^. While our data reflect acute modulation in healthy young adults, the consistency of *θ* enhancement across two independent experiments suggests that rhythmic oral input may transiently engage the same sensorimotor pathways that support long-term plasticity. In this sense, chewing may act both as an immediate entrainer and as a behavioral tool that preserves network flexibility in hippocampal–prefrontal circuitry.

### Integration with MoBI and embodied cognition frameworks

Recent Mobile Brain/Body Imaging (MoBI) frameworks emphasize that cognition unfolds within continuous sensorimotor activity and that bodily rhythms provide temporal scaffolds for cortical computation^19^. Our findings align closely with this view. Converging evidence across independent cognitive contexts—including our complementary post-chewing dataset and prior chewing paradigms—demonstrates that rhythmic mastication consistently engages a frontocentral *θ* network. *θ* increases over AFz, Fz, and FCz emerge in the same frequency range and direction across studies, despite differences in task timing, stimulus properties, and preprocessing pipelines. This reproducibility underscores that chewing is not an epiphenomenon of one task but a general sensorimotor modulator of executive networks.

### Summary and implications

This study identifies chewing as a reliable and replicable modulator of cortical *θ* oscillations and cognitive control. Two independent experiments—one with chewing during task performance and one with chewing prior to task onset—produced convergent behavioral improvements and overlapping electrophysiological signatures. This replication across experimental designs strengthens causal interpretation and situates mastication within a broader class of bodily rhythms that interact with cortical computation through oscillatory entrainment.

These results advance the understanding of sensorimotor contributions to cognition and suggest that preserving or restoring masticatory function may benefit executive performance. Beyond cognitive enhancement, the modulation of frontoparietal *θ* may also intersect with motor control and balance, domains in which masticatory dysfunction has measurable consequences. Chewing emerges as an accessible, non-invasive, and ecologically valid means of engaging sensorimotor pathways that support large-scale neural coordination.

### Limitations and future directions

Several limitations should be noted. EEG lacks spatial resolution to measure thalamic, brainstem, or hippocampal activity directly, and future studies should incorporate MEG, EEG–fMRI, or invasive recordings in animal models. Direct manipulations of trigeminal input (e.g., local anesthesia or nerve blocks) and neuromodulatory systems will help disentangle afferent versus gain-related mechanisms. Our sample comprised healthy young adults; whether similar mechanisms operate in older individuals or in those with masticatory dysfunction remains to be determined. Finally, computational modeling approaches such as Dynamic Causal Modeling (DCM) could formally test directionality in thalamocortical and frontoparietal interactions during chewing.

## Conclusion

Across two independent experiments, we demonstrate that chewing enhances cognitive performance and modulates frontocentral *θ* dynamics through rhythmic entrainment. These findings support an embodied account of cognition in which peripheral sensorimotor rhythms shape cortical processing in real time and facilitate large-scale network coordination. Mastication thus emerges as a powerful yet underappreciated modulator of executive function, with implications spanning neuroscience, dentistry, and motor control.

## Methods

### Participants

Forty-six healthy right-handed adults participated. None reported falls in the previous year, symptoms of temporomandibular disorders (per DC-TMD^41^), or use of psychiatric or mood-altering medication. Participants were randomly assigned to a chewing (case) or non-chewing (control) group. The case group performed the second block of the task while chewing a specially formulated gum; controls performed both blocks without chewing.

### Behavioral task

A visuospatial 2-back task^29^ was used. A small square appeared for 200 ms in a 3 × 3 grid with a 2000 ms inter-stimulus interval. Participants pressed a button when the position matched the item two trials earlier (25% targets). Stimuli were presented within 5° of visual angle and consecutive targets were prevented.

### Statistical analysis

All analyses were performed in MATLAB R2024b. Normality was assessed using the Kolmogorov–Smirnov test; because several variables violated this assumption, non-parametric statistics were used throughout. Behavioral performance (RT, ACC, and IES^42^) was compared across blocks using paired Wilcoxon signed-rank tests, whereas between-group comparisons (chew vs. control) used Mann–Whitney U tests. Effect sizes were reported as Cliff’s delta (*δ*). *θ*-band power (4–7 Hz) was averaged within two predefined temporal windows (early: 100–900 ms; late: 900–1300 ms) and compared across conditions using paired Wilcoxon signed-rank tests. For PAC analyses, Modulation Index (MI) values were tested against zero using one-sample Wilcoxon tests, preferred phases were evaluated using the Rayleigh test for circular non-uniformity, and the proportion of participants with significant MI values was assessed with a binomial test. Robustness of MI estimates was further evaluated by comparing blocks with different recording lengths (Mann–Whitney), applying equal-*N* resampling (paired shift test; Pearson correlation)^43^, generating surrogate-based null distributions (1,000 time-shift permutations; *z*-scored MI), and removing the aperiodic 1/ *f* component using the FOOOF model followed by a Wilcoxon test on the *θ*-band residuals. All statistical tests were restricted to predefined ROIs, frequencies, and time windows based on a priori hypotheses; therefore, no post-hoc correction was applied. Statistical significance was defined as *p <* 0.05 (two-tailed).

### EEG acquisition

Stimuli were presented with PsychoPy^44^ while EEG/sEMG was recorded using a BioSemi ActiveTwo 64+8 channel system at 1 kHz. Electrodes followed the 10–20 layout. Recordings were conducted in an electrically shielded chamber.

#### Pipeline validation

We evaluated three candidate preprocessing approaches: (1) ICanClean^20^ with a pseudo-reference, (2) artifact subspace reconstruction (ASR)^26^ calibrated on a 3-min resting-state baseline, and (3) a 3–20 Hz bandstop filter (order 398), as well as combined pipelines. Spectral structure and time–frequency representations were compared against a traditionally cleaned dataset (1 Hz high-pass, ICA, interpolation). The combined ICanClean+ASR method (M4) provided the best artifact suppression under chewing conditions (see Supplementary Fig. S3).

#### EEG preprocessing

Continuous EEG was high-pass filtered at 0.1 Hz, downsampled to 256 Hz, and denoised using Zapline-plus^45^. ASR^26^ was applied using each participant’s cleaned resting-state data as a calibration template. Residual artifacts were removed via ICA using runica^46^. Components reflecting eye movements, muscle activity or line noise were rejected. For chewing blocks, a pre-ICA cleaning step using ICanClean^20^ (threshold *ρ* = 0.77) and a 1–20 Hz bandstop filter was applied.

#### Surface EMG

Masseter activity was recorded using monopolar BioSemi electrodes and converted to bipolar signals. EMG was bandpass filtered (20–400 Hz), rectified and normalized. The envelope was extracted using a 100 ms RMS window. Chewing cycles were detected via adaptive amplitude thresholding and a minimum 0.5 s inter-peak interval. Chewing frequency was computed as the number of detected peaks divided by the recording duration.

#### Validation of EEG signal integrity

To verify that chewing-related EMG did not contaminate EEG *θ* activity, we quantified PAC between EMG phase (1–2 Hz) and EEG amplitude (4–40 Hz). *θ*-range PAC (4–8 Hz) was minimal (median MI_*θ*_ = 1.66 × 10^−4^) and substantially lower than beta or high-frequency PAC. PAC did not correlate with EMG power (*p >* 0.21). Topographically, EMG-driven PAC (20–40 Hz) was localized to temporal sites, whereas *θ*-band PAC showed a flat distribution with no enhancement in the frontocentral ROI. This confirms that frontocentral *θ* activity reflects cortical rather than muscular sources.

#### Time–frequency analysis

Time–frequency decompositions were computed using Morlet wavelets (4–13 cycles) across 400 frequencies (1–40 Hz; 0.1 Hz resolution). Power was averaged across trials and converted to decibels relative to a pre-stimulus baseline.

#### Frontocentral region of interest

All time–frequency and PAC analyses focused on a predefined frontocentral region of interest (ROI). This ROI comprised electrodes AFz, F1, Fz, F2, FC1, FCz and FC2, which were pooled by averaging power or amplitude across channels for each subject and condition.

The choice of this ROI was guided by three considerations. First, in our previous work using a similar chewing paradigm, chewing-related increases in *θ* power and connectivity were maximal over frontocentral sites^**?**^ (REF: Paper 1). Second, frontocentral *θ* activity is consistently implicated in visuospatial working memory and oddball processing in the EEG literature, often peaking around midline frontal and frontocentral electrodes. Third, inspection of grand-average topographies in the present dataset confirmed that the main *θ* effects fell within this predefined cluster, without extending into temporal regions typically dominated by EMG activity.

To avoid circularity, the ROI was defined *a priori* and was not adjusted based on any between-condition contrast. All statistical tests on *θ*-band power and PAC were therefore restricted to this frontocentral ROI.

#### Phase–amplitude coupling and Modulation Index calculation

PAC between chewing rhythm and cortical *θ* amplitude was quantified using the Modulation Index (MI)^30^. EMG phase (1–2 Hz) was obtained via the Hilbert transform. *θ*-band amplitude (4–8 Hz) was extracted from band-passed EEG and averaged over a predefined frontocentral ROI (AFz, F1, Fz, F2, FC1, FCz, FC2). Amplitude values were binned into 18 phase bins to form a distribution *P*. MI was computed as:

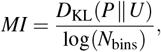

with *N*_bins_ = 18. Preferred phase was derived from the first circular moment. PAC was computed on continuous data to avoid edge artifacts.

## Ethics

The study was approved by the ethics committee of the School of Dentistry, Universidad de Valparaíso (POSTG-06-22). Procedures complied with the Declaration of Helsinki. Written informed consent was obtained from all participants.

## Acknowledgements

The authors acknowledge ANID, Chile.

Beca Doctorado Nacional 21220968 (SE).

FONDECYT 1241965, Exploracion 13240064, and AC3E CIA250006 (WE).

## Author contributions statement

S.E. & W.E. conceived the experiment, S.E. conducted the experiment, & S.E. analysed the results. D.M. & L.M. formulated the chewing gum. All authors reviewed the manuscript.

## Additional information

### Data avaibility

All data, raw and preprocessed, and codes for reproducing figures under DOI 10.5281/zenodo.16876077

## Supplementary Figures

**Supplementary Figure S1.**
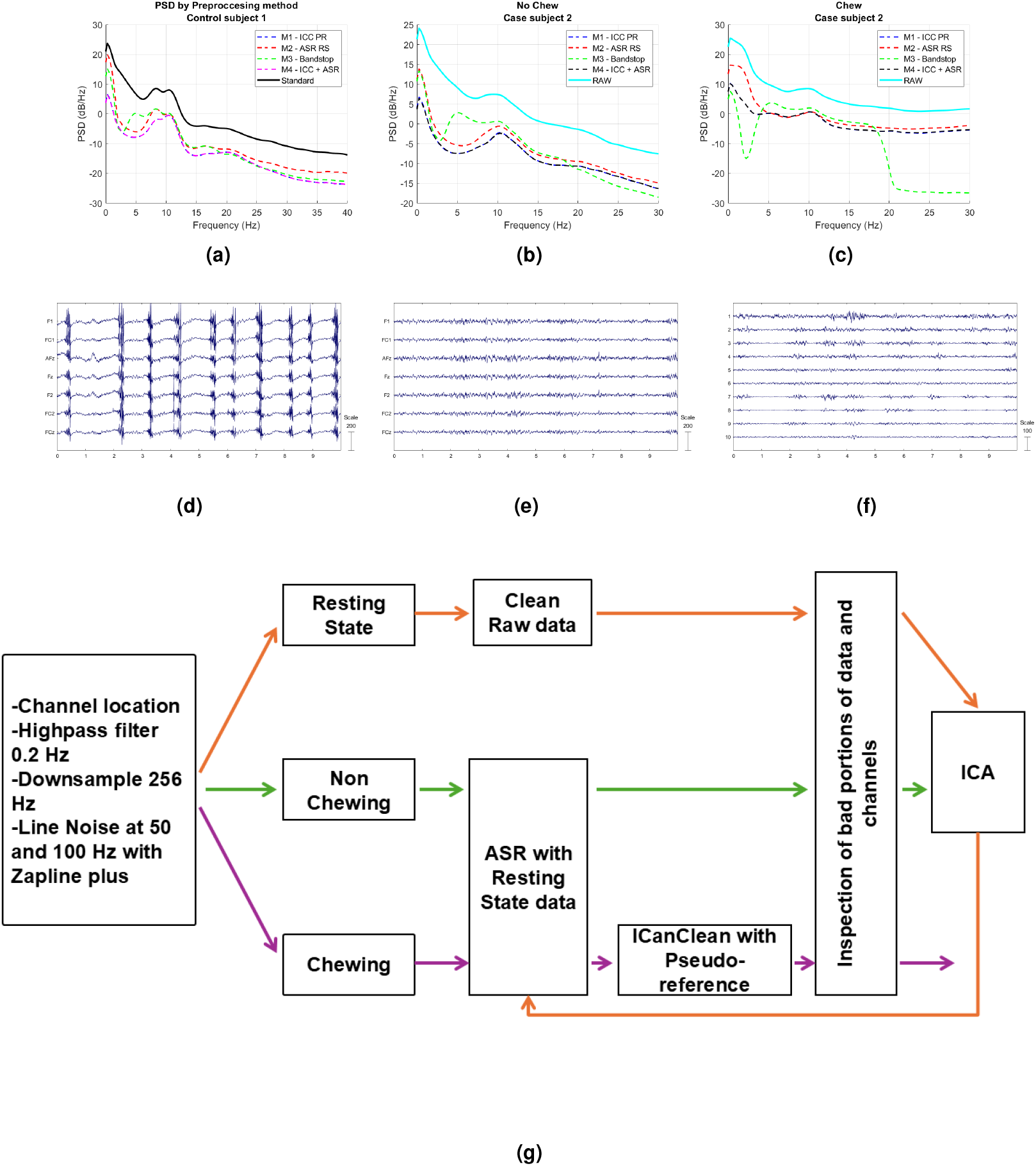
Validation of the preprocessing pipeline: a Power spectral density comparison of the standard method and alternative methods; b PSD of different methods without chewing; c PSD of different methods with chewing; d Raw EEG data of the subject; e Cleaned data at the same scale and time window; f Independent components obtained after cleaning; g Preprocessing pipeline.

**Supplementary Figure S2.**
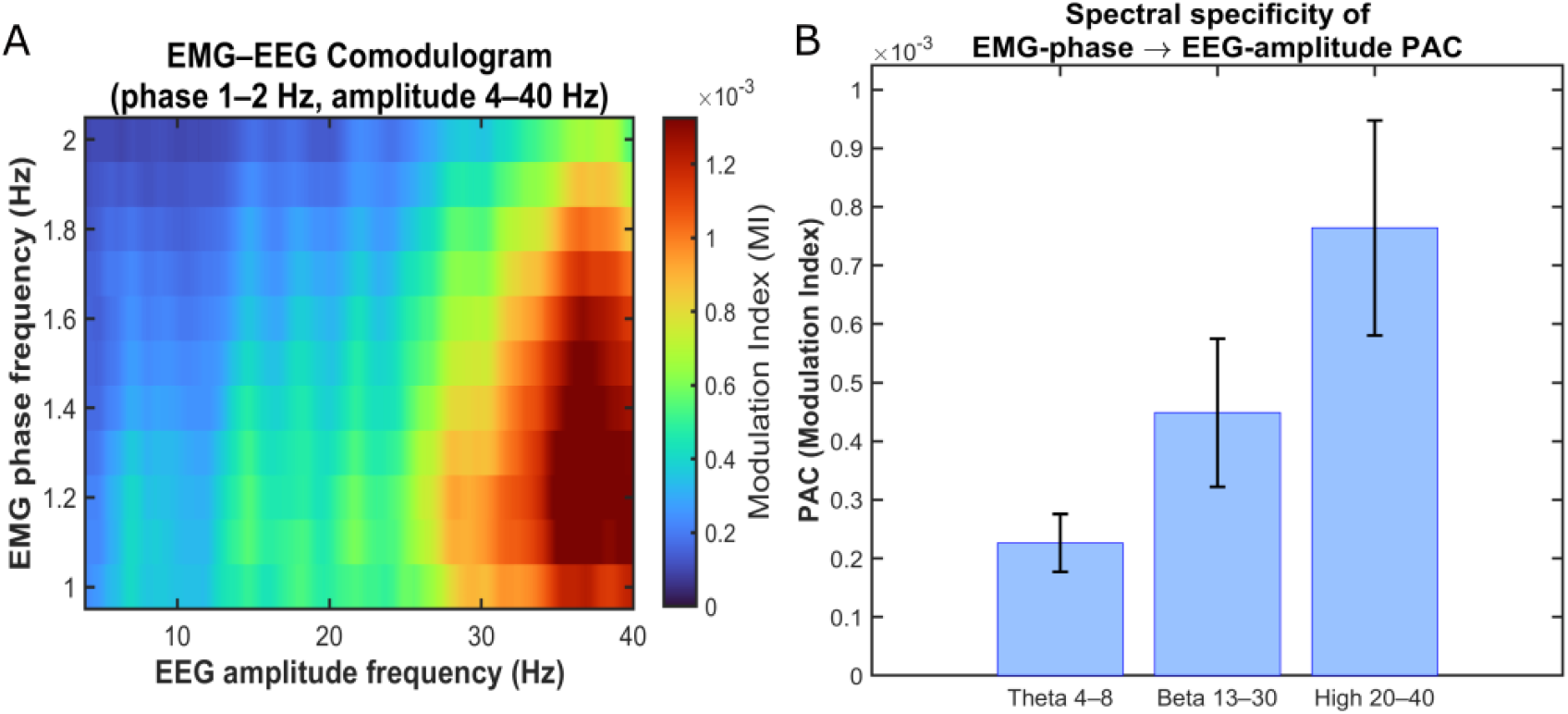
Chewing-related EMG does not selectively drive frontocentral theta. (**A**) Group-level comodulogram of phase–amplitude coupling (PAC) between the masticatory EMG phase (1–2 Hz) and EEG amplitude (4–40 Hz), averaged over the frontocentral ROI. PAC values are overall small in the theta range (4–8 Hz) and increase gradually towards higher frequencies, peaking above 20 Hz, consistent with the typical high-frequency spectrum of facial EMG. (**B**) Mean PAC (modulation index) for theta (4–8 Hz), beta (13–30 Hz), and high frequencies (20–40 Hz) bands (error bars: SEM across participants, *N* = 31). Theta-band PAC is weak and not selectively enhanced compared with higher-frequency ranges, indicating that rhythmic chewing-related EMG does not account for the frontocentral theta activity analyzed in the main results.

**Supplementary Figure S3.**
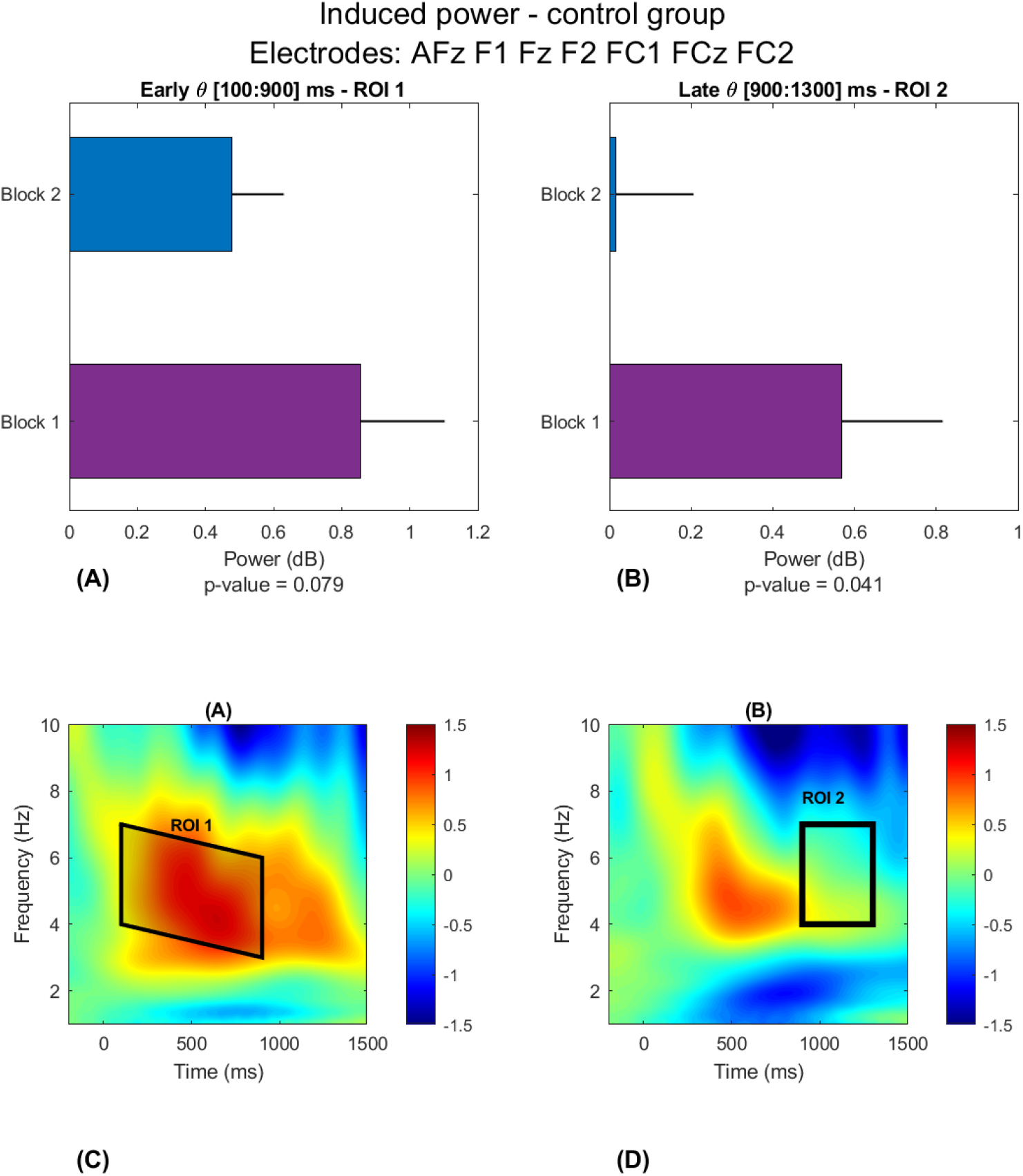
Control group showed a contrary behavior of the *θ* induced-power. **(A)**, showed no statistically significant differences between blocks 1 and 2 in the ROI 1, just as Figure 2 (Panels C and D), while **(B)** shows a decrease in *θ* contrary to Figure 2 (Panels E and F) were recovers and increase the induced power.

